# Odorant Mixture Separation in *Drosophila* Early Olfactory System

**DOI:** 10.1101/2022.09.20.508558

**Authors:** Aurel A. Lazar, Tingkai Liu, Chung-Heng Yeh, Yiyin Zhou

## Abstract

Natural odorant scenes are complex landscapes comprising mixtures of volatile compounds. It was previously proposed that the Antennal Lobe circuit recovers the odorant identity in a concentration-invariant manner via divisive normalization of Local Neurons. It remains unclear, however, how identities of odorant components in a *mixture* is represented or recovered in the fruit fly early olfactory pathway. In the current work, we take a different approach from the traditional steady-state analyses that classify odorant mixture encoding into configural vs. elemental schemes. Instead, we focus on the spatio-temporal responses of the early olfactory pathway at the levels of the Antennal Lobe and the Mushroom Body, and formulate the odorant demixing problem as a blind source separation problem - where the identities of each individual odorant component and their corresponding concentration waveforms are recovered from the spatio-temporal PSTH of Olfactory Sensory Neurons (OSNs), Projection Neurons (PNs), and Kenyon Cells (KCs) respectively. Building upon previous models of the Antenna and the Antennal Lobe, we advanced a feedback divisive normalization architecture of the Mushroom Body Calyx circuit comprised of PN, KC and the giant Anterior Paired Lateral (APL) neuron. We demonstrate that the PN-KC-APL circuit produces a high dimensional representation of odorant mixture with robust sparsity, and results in greater odorant demixing performance than the PN responses.

## 1 Introduction

Neural systems implement sophisticated internal representations of sensory information for performing complex computation and cognitive functions. For insects with ecological niche emphasizing olfactory navigation, robust representation of odorant mixture is achieved by an evolutionarily conserved neural pathway. In particular, we focus on the first three layers of the fruit fly early olfactory pathway that is comprised of the Antenna, the Antennal Lobe and the Calyx of the Mushroom Body as shown in **Fig.**1, and sought to explore the odorant mixture processing capabilities of each of the three layers.

**Figure 1:**
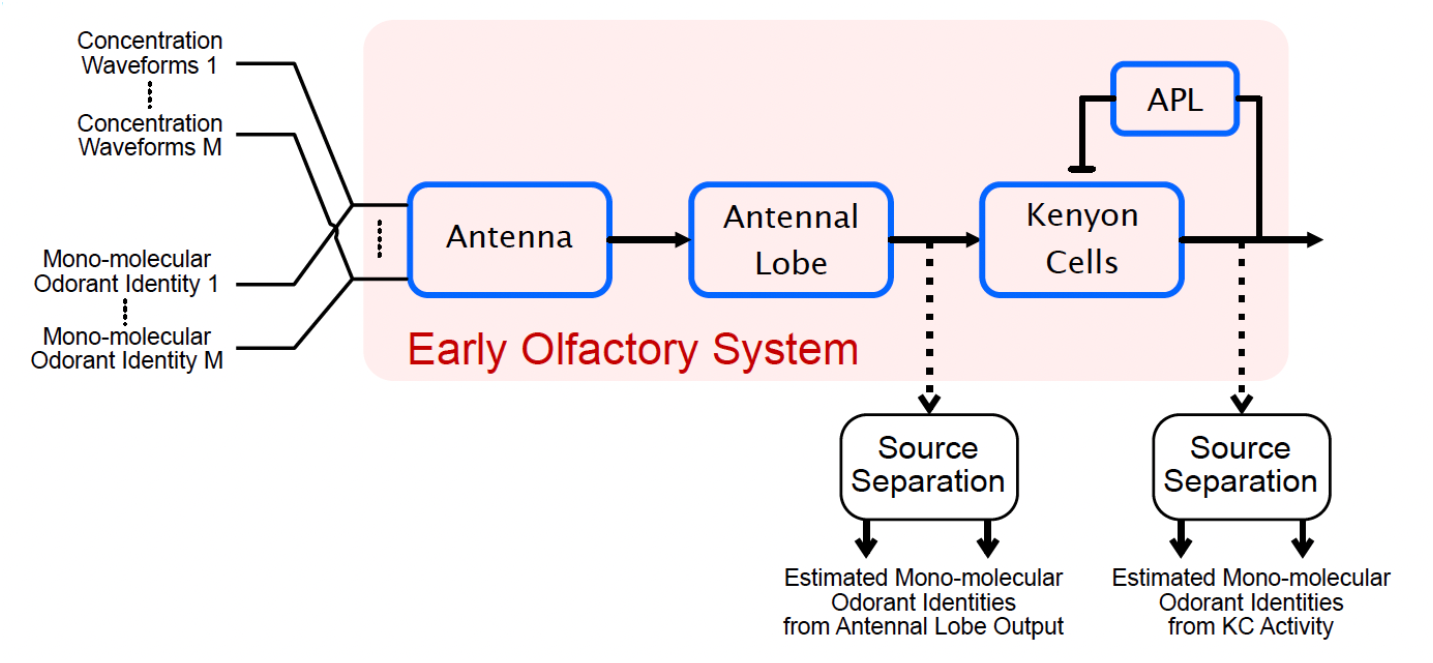
The early olfactory pathway with odorant mixture input, and the odorant demixing problem at the level of Antennal Lobe (Projection Neurons) and the Calyx of the Mushroom Body (Kenyon Cells).

It has previously been proposed that the Antenna encodes odorant identity and concentration waveform via multiplicative coupling [1], and the Antennal Lobe separates the odorant identity and concentration waveform via an ON-OFF odorant objective identity recovery processor implemented by the Local Neuron circuits [2]. However, the discussion of odorant identity recovery in the Antennal Lobe only pertains to mono-molecular odorants, and it remains unclear how odorant mixtures are represented in the early olfactory pathway.

In the current work, we advanced a feedback Divisive Normalization model of the PN-KC-APL circuit consisting of biophysical neurons and synapse models. The proposed PN-KC-APL circuit encodes odorant stimuli, pure and mixture alike, into time-dependent high dimensional spatio-temporal KC responses that show robust sparsity across time, odorant identities and concentration waveforms. Furthermore, by formulating odorant mixture processing as a blind source separation problem (see **Fig.**1), we show that the KC responses enable better identifications of odorant components in a mixture. As opposed to the dichotomy of Elemental vs. Configural encoding previously discussed in mixture processing [3, 4, 5], our focus on the spatio-temporal responses of the OSNs, the PNs and the KCs enabled us to show that odorant component identities can be recovered from all layers of the olfactory pathway to varying degrees, demonstrating that all odorant representations in the olfactory pathway are elemental.

Finally, we show that the odorant demixing capability of the KC responses depends on the expansion-normalization circuit architecture, which leads to a high dimensional Voronoi partitioning of the odorant space by the KC representation. This geometric perspective of the KC representation of odorants shows that odorant representation by KCs is locally robust and globally sensitive to changes in odorant mixture compositions, and explains why odorant demixing is easier at the level of the KCs.

## 2 Odorant Mixture Representations in the Early Olfactory System

### 2.1 Modeling the Space of Odorant Mixtures

We extended the model of pure odorant input [1] to odorant mixture by assuming that the odorant mixture drives the olfactory receptor in a linear weighted sum fashion, thereby extending the synthetic odorant DB proposed in [1]:

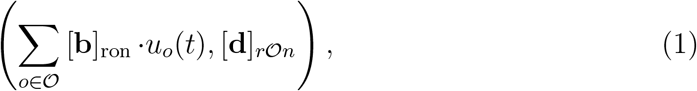

where 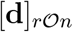 describes the effective dissociation rate of the mixture, which is defined as the weighted mean of dissociation rates for each individual odorant-receptor pair

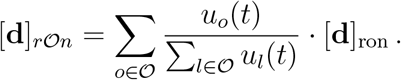

We showed that this simple linear mixture model reproduces experimental observation of competitive binding in olfactory sensory neurons.

### 2.2 Modeling the Expansion/Normalization Architecture of the PN-KC-APL Circuit

We then introduce the feedback divisive normalization architecture of the PN-KC-APL circuit (see **Fig.**2), its parameterizations and their influence on the output Kenyon Cell responses.

**Figure 2:**
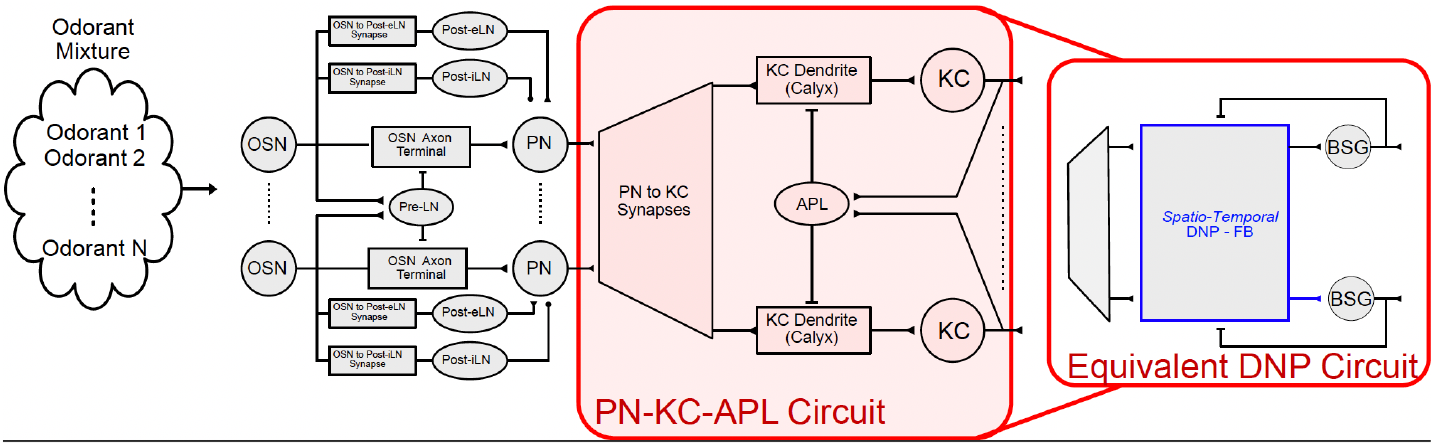
Model of odorant mixture processing in the early olfactory system and the PN-KC-APL circuit architecture.

First described in [6], over 150 years of studies on MB provide detailed, numerous anatomical and physiological properties of the KC circuitry. The estimated number of KCs is ≈ 2, 000 [7]. KCs receive excitatory inputs from projection neurons (PN) in the antennal lobe [8, 9]. Each PN innervates one of 50 sub-neuropil structures, called a glomerulus [10, 11, 12]. PNs connect randomly to KCs [8]. On average, each KC is driven by 6.2 PNs with some biases [8, 9]. Note that KCs also receive input from sensory modalities other than the olfactory system [13, 14, 15]. These additional inputs, however, are not the focus of this work. A key feature of the PN-KC-APL circuit is that all KCs excite a giant GABAergic anterior paired lateral (APL) neuron, which reciprocally inhibits all KCs [16, 17, 18]. While detailed connectomic data of the adult fly brain reveals additional complexity of KC-APL connection [19], the reciprocity is observed in the larva fly brain [20]. For Simplicity, we assume a global feedback inhibitory APL to KC connection. Regarding the physiological characteristics of the APL neuron, its putative analog in the locust (the giant GABAergic neuron (GGN) [21, 22]) is non-spiking, suggesting that the APL neuron is likely a graded potential neuron [23].

Physiologically, it has been reported that the PN response is dense for both monomolecular odorants and mixtures [24, 25], while the KC responses are 5 ~ 10% sparse (only 5–10% KCs react to an odorant stimulus at a time) [26, 27, 28, 18]. This sparsity has been attributed to the feedback inhibition from APL to KC [21, 29, 30, 26, 31, 32]. While the exact APL to KC inhibition mechanism in the Mushroom Body Calyx is unclear, we applied the differential Divisive Normalization Processor model proposed in [2] to describe the PN-KC-APL interaction in the KC Dendritic Tree. Therefore, the PN-KC-APL circuit (shown in **Fig.**2) is comprised of a PN-to-KC expansion circuit and a KC-APL feedback normalization circuit.

## 3 Algorithms for Odorant Demixing

Assume that the PN/KC responses to pure odorants is known, the odorant demixing problem seeks to evaluate the mixture processing of the olfactory pathway by comparing the abilities to determine odorant component identities from the PN/KC spatio-temporal mixture responses (spike rates) under the same concentration amplitude.

### Assumption

We assume that the steady state response (spike rate) to pure odorants of constant concentration *u* is known. We denote the vector of steady state response vectors (PSTH) to an odorant *o* by **r**_*o*_(*u*), *o* = 1,…, *N_O_*, for a total of *N_O_* = 110 pure odorants. We define the dictionary of pure odorant responses as

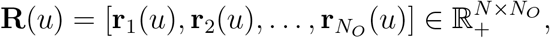

where *N* is the appropriate dimension of the population responses, e.g., *N* = 50 for PNs and *N* = 2000 for KCs. While PN response is significantly less concentration dependent than the OSN response [2], the dictionary **R** is parameterized by concentration *u* to account for imperfect concentration-invariance of PN response.

### Problem Definition

We denote the response to an odorant mixture as **x**(*t*). The demixing problem is formulated as: given **R**(*u*), identify the odorant components of the mixture from **x**(*t*).

#### • Algorithm

Minimize the Frobenius norm

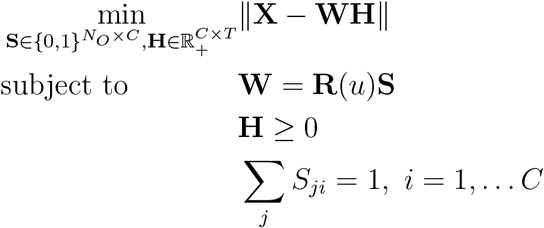

where 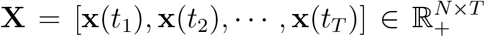 is the mixture representation of PNs or KCs sampled at different time instances *T* = {*t*_1_, *t*_2_,…, *t_T_*}. We assume that the mixture has *C* odorant components, and **S** has *C* columns - each column vector is an indicator vector with only 1 non-zero entry. Therefore, the constraint **W** = **R**(*u*)**S** restricts the columns of **W** to be a subset of the columns of **R**(*u*). **H** is a non-negative matrix.

According to the model of the odorant space, an odorant is defined by its identity and concentration. The identity is a time-invariant concept whereas concentration varies over time. Therefore, the goal of the minimization above is to factorize **X** into **W** and **H**, where **W** is time-invariant and represents the identities of component odorants and **H** represents the “concentration waveform” of each mixture component in each row. At any single time instance, there can be many choices to combine different identities and concentrations together to obtain the response at that time instance, and thereby the choice is ambiguous. The optimization problem explores the constancy of the identities of mixture components across time to reduce this ambiguity. Thus, we are basically assuming that [**X**]_*ji*_, the response of neuron (PN or KC) *j* at time *t_i_*, can be linearly decomposed as [**X**]_*ji*_ = [**r**_*o*_1__]_*j*_ · [**H**]_1*i*_ + [**r**_*O*_2__]_*j*_ · [**H**]_2*i*_ +…+ [**r***_o_C__*]_*j*_ · [**H**]_*Ci*_, where *o*_1_, *o*_2_,…, *o_C_*, are the C indices of columns of **R**(*u*).

#### • Evaluation

To evaluate accuracy of demixing a binary odorant mixture, we define a score of 1 if both components are identified correctly, 0.5 if only one component is identified correctly and 0 if neither is identified. The score is averaged over all pairs of mixture components.

## 4 Results

### 4.1 Expansion-Normalization Produces KC Responses with Robust Sparsity

We showed that, by choosing parameters of the circuit in **Fig.**2 to enable strong APL inhibition of KC Dendrite (**Fig.3**(B)), the KC response is robustly sparse across odorant identities (**Fig.**3(A)), concentrations (**Fig.**3(A)) and time (**Fig.**3(D)). We also showed in **Fig.**3(C) that the sparsity level of the KC output is independent of the number of KCs, and can be entirely controlled by changing the spiking threshold of the KC Biophysical Spike Generator (BSG) and the number of PNs projecting onto the same KC (PN-KC fan-in). Given the tight control over the sparsity of the KC response, we therefore consider different PN-KC-APL circuit models that are parameterized by 1) the number of KCs and 2) the KC response sparsity level for subsequent comparative evaluations of mixture processing.

**Figure 3:**
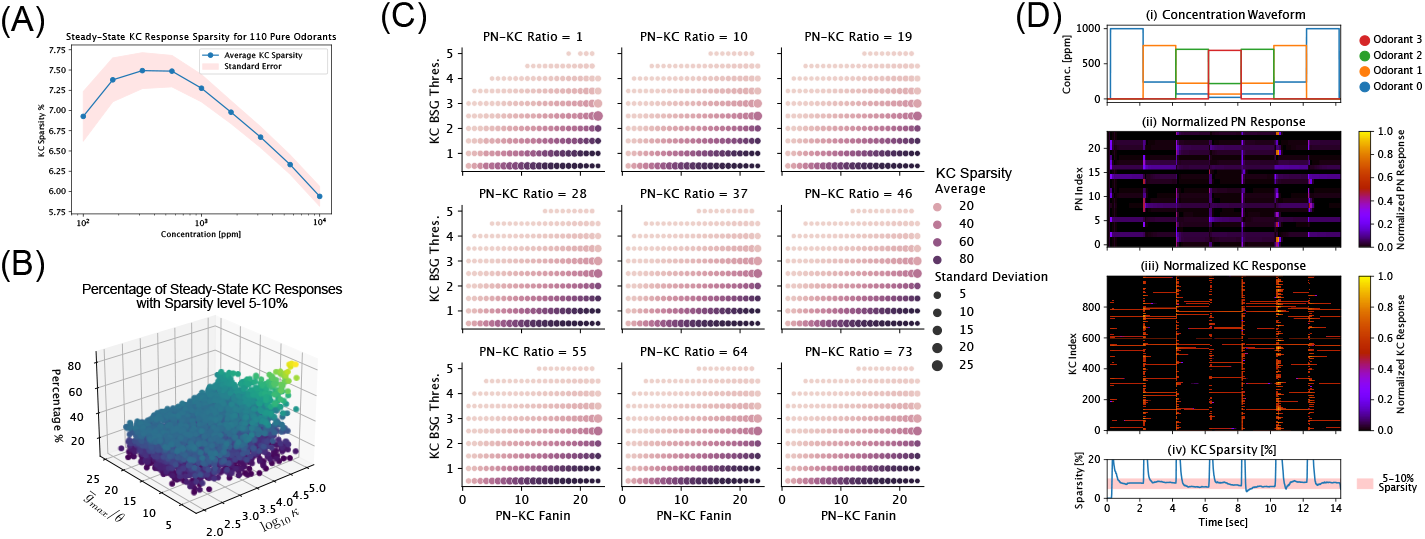
Parametrization of PN-KC-APL circuit enables robust sparsity of KC responses across odorant identities, concentration and time. (A) KC sparsity across all 110 pure odorants with known affinity values [1, 33] and concentrations spanning 4 orders of magnitudes. Blue solid curve shows the average KC sparsity at each concentration level, light blue band around the mean value shows +/− 1 standard error of KC sparsity level. (B) KC sparsity robustness vs. parameters of the KC DNP model, for PN-KC-APL circuit with 1000 KCs and 7 PN-KC fanin. Each dot represents a parametrization of the KC DNP model, and the dot is color-coded by the percentage of the KC steady-state responses with sparsity within the 5-10% range. Note that the most robustly sparse parametrization (top right corner, color-coded in yellow) is when *κ* is large, indicating strong APL inhibition of KC Dendrite. (C) Average KC sparsity across concentration and odorant identities compared against number of KCs, PN-KC fan-in ratio and KC BSG threshold. The average KC sparsity is determined by the PN-KC fan-in and KC BSG threshold, and is independent of the number of KCs as all subfigures have similar KC average sparsity for a given (PN-KC fan-in, KC BSG threshold) pair. (D) Example I/O and KC sparsity of binary mixture input. (1st row) Concentration waveforms of two odorant components in the mixture. (2nd row) PN spatial temporal spike rate (normalized to between [0,1]). (3rd row) KC spatial temporal spike rate (normalized to between [0,1]). (4th row) KC sparsity across time.

### 4.2 Expansion-Normalization Promotes Foreground/Background Mixture Separation in KC Responses

Based on the methodology described in the previous section, we compared the accuracy of demixing using the mixture representation before and after the PN-KC-APL circuit (**Fig.**1).

With a total of 110 pure odorant components, 5995 binary odorant mixtures can be constructed. Due to computational constraints, we randomly selected 100/5995 binary mixtures, each driven at 7 different concentration levels *u* at 9 different mixture ratios *θ*, resulting in a total of 6, 400 experiments for each circuit configuration.

The same 6, 400 experiments are repeated for 4 different PN-to-KC expansion ratios [1, 5, 10, 40] (note that the PN-to-KC expansion ratio in the 1st instar larval Mushroom Body is ≈ 5) each parameterized to produce KC steady-state responses at 3 different sparsity levels (5 ~ 10%, 20 ~ 30%, 40 ~ 50%). In total, without changing the Antenna and Antennal Lobe circuits, 12 PN-KC-APL circuits are simulated, each with 6, 400 different binary mixture inputs. Each of the experiment is simulated for 5 seconds (simulation time) at 0.1 millisecond temporal resolution, with concentration waveforms having temporal supports of [0.2, 4.8] seconds. The resulting PN and KC mixture spatio-temporal responses are sampled at 50 millisecond intervals, resulting in 100 discretized time steps.

From each source separation result (a total of 32, 000 for PN and KC, respectively), we compute the identification accuracy *acc^K^,acc^P^*, where *acc^K^* ∈ [0, 1] is the accruacy of identifying odorant components from mixture KC response as defined above, and resp. *acc^P^* from PN response. Shown in **Fig.**4(main figure) is the average difference in identification accuracy *acc^K^* – *acc^P^*, where each rectangle shows the differences across concentration/mixture ratios (horizontal axis) and overall mixture concentrations (vertical axis). In **Fig.**4(inset), the difference in identification accuracy is further averaged across concentrations and mixture ratios, resulting in a scalar value for each of the 12 circuit configuration. The differences are color-coded on a Blue(negative)-to-Red(positive) scale, with Red corresponding to higher KC identification accuracy than PN, and vice versa. As shown in **Fig.**4(inset), KC outperforms PN in identification accuracy for high expansion ratio level and with sparser KC responses (lower sparisty level), this suggests that the odorant mixture separation at the level of KC requires the Expansion-and-Sparsification in the PN-KC-APL circuit. In summary, our results showed that, to achieve better KC odorant demixing performance, both a high PN-KC expansion ratio and strong APL inhibition of the KC Dendrite (leading to low KC sparsity level) are required.

**Figure 4:**
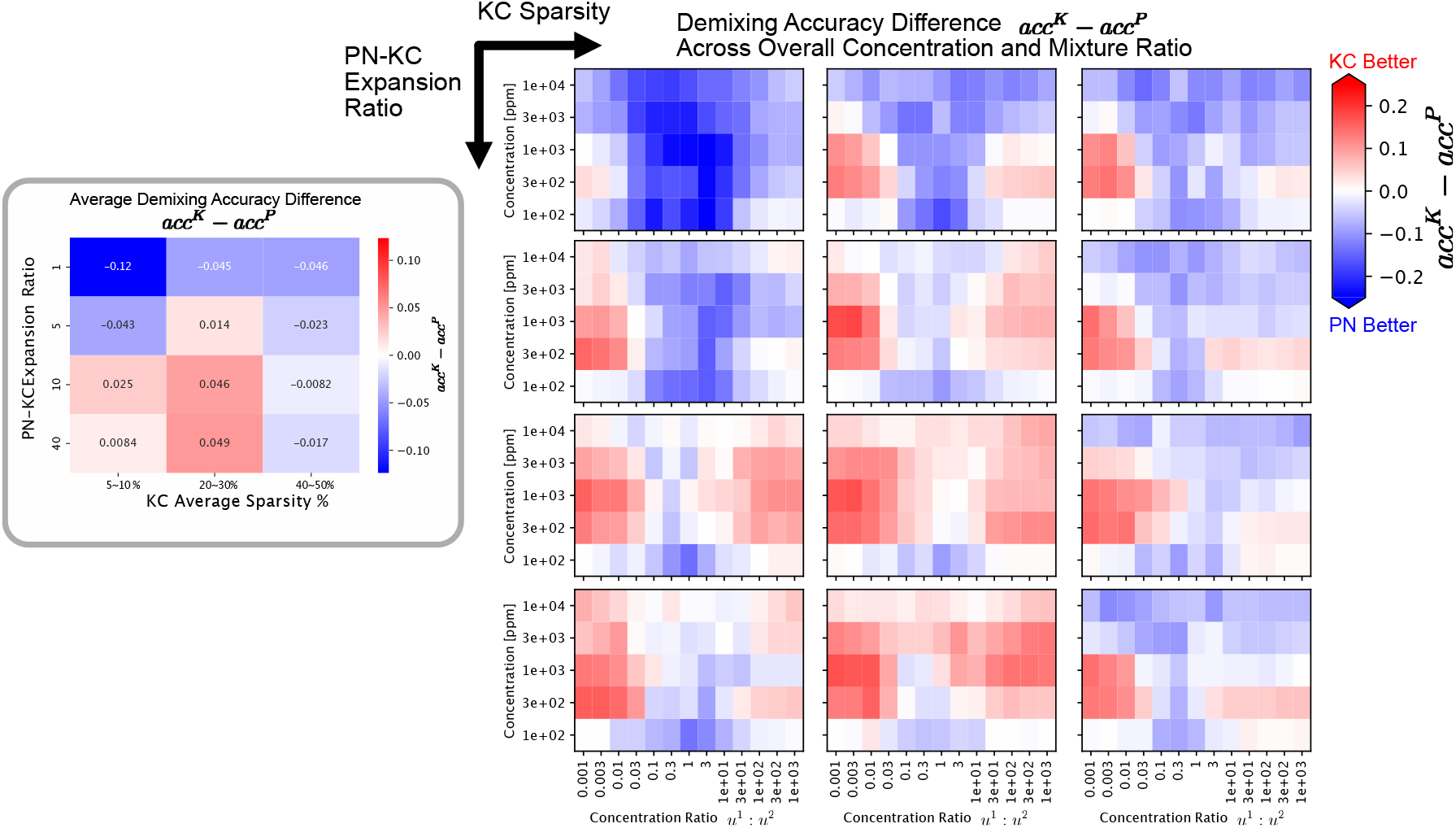
Binary mixture demxing accuracy comparison between PN *acc^P^* and KC *acc^K^*. Heatmaps are color-coded by the difference in recovery accuracy *acc^K^* – *acc^P^*, where higher (Red) indicates better recovery at the KC level than the PN level, lower (Blue) indicates better recovery at the PN level than the KC level. (Inset) Average difference in demixing accuracy *acc^K^* – *acc^P^* across mixture identities, concentration and ratio for increasing PN-KC Expansion Ratios (from top to bottom) and increasing KC average sparsrity levels (from left to right). (Main Figure) Average difference in demixing accuracy *acc^K^* – *acc^P^* across mixture identities, plotted for each mixture concentration and ratio. Each of the 12 subfigures correspond to a circuit architecture with specific corresponding (PN-KC Expansion Ratios, KC Sparsity) as in the inset. The inset shows KC has higher demixing accuracy than PN for higher expansion ratio (expansion) and low sparsity (sparsification), while the main figure show that the KC demixing accuracy is higher at very high/low concentration ratios (strong foreground and weak backround) and lower at intermediate concentration ratios.

## Acknowledgments

The research reported here was supported by AFOSR under grant #FA9550-16-1-0410, DARPA under contract #HR0011-19-9-0035 and NSF under grant #2024607.

